# S2F-Agent: Harnessing sequence-to-function models for verifiable genome interpretation

**DOI:** 10.64898/2026.05.13.724757

**Authors:** Jiaqi Li, Ti Qin, Jingyu Li, Zhiwei Bao

## Abstract

Sequence-to-function (S2F) models offer a revolutionary paradigm for genotype-phenotype mapping, yet their broader application is bottlenecked by the need for reliable orchestration and interpretation across a fragmented model ecosystem. While general-purpose language models can automate scientific workflows, they are not inherently grounded in the model-specific execution constraints required for robust S2F analysis. Here, we present S2F-Agent, a human-in-the-loop framework designed for the verifiable orchestration of the heterogeneous S2F ecosystems. The framework employs a contract-based harness to bridge model-specific capabilities (Skills) and model-agnostic biological objectives (Playbooks), seamlessly translating free-form biological requests into reliable execution and rigorous downstream interpretation. Evaluated on a benchmark of 54 query cases derived from published S2F workflows, S2F-Agent systematically outperformed general-purpose LLMs, demonstrating superior reliability accuracy in routing, groundedness, and end-to-end task execution success. We further demonstrate the robustness and scalability of S2F-Agent across model adaptation, variant interpretation, genome-scale functional profiling and personal-genome analysis. First, the agent autonomously adapts a genomic foundation model to quantitative chromatin profiles, resolving sequence features associated with primed and active regulatory states. Second, integrating multi-perspective variant effect predictions prioritized 42 high-priority candidate variants among CAD-associated variants (>16,000), and identified tissue-resolved regulatory mechanisms including the hepatic SORT1 axis. Third, genome-scale profiling of multiple traits GWAS atlas variants (>250,000) revealed pervasive context dependence in molecular consequences and regulatory architecture, highlighting the analytical focus toward fine-grained, tissue-specific regulatory variants. Finally, evidence-gated analysis of personal genomes expanded functional hypothesis generation beyond clinically annotated variants to thousands of prioritized candidates per individual while imposing explicit evidence-dependent boundaries on clinical claims. Collectively, these results establish S2F-Agent as a general framework for converting heterogeneous sequence-to-function capabilities into verifiable, scalable, and evidence-aware genomic analyses. By bridging the chasm between LLMs, specialized S2F ecosystems and rigorous genomic science, this framework democratizes the S2F paradigm for unlocking the full potential of these advanced models in real-world discoveries.

## Introduction

A central challenge in modern biology is to understand how DNA sequence encodes molecular function, particularly in the non-coding genome where most trait- and disease-associated variants reside^1^. Sequence-to-function (S2F) models have substantially advanced this problem by learning sequence features that predict diverse cell-type-specific molecular phenotypes, including chromatin accessibility, transcription, splicing, and evolutionary constraint, thereby enabling the identification of cis-regulatory elements to the interpretation of non-coding variants^2^. However, with the explosive growth of the S2F ecosystem, the critical bottleneck is shifting from model capability to reliable orchestration and interpretation across heterogeneous models. Different models operate across distinct sequence representations, genomic contexts, and context-specific functional outputs, while actionable biological insights often require integration with heterogeneous predictions and orthogonal evidence before they can support robust claims. Consequently, increasingly sophisticated S2F analyses remain highly dependent on laborious expert-driven workflow design, parameterization, and cross-model integration. Overcoming this gap requires the reliable orchestration of automated workflows that can scale manual analyses, ultimately making S2F capabilities reproducible, scalable, and broadly accessible to the scientific community.

While recent agentic systems have demonstrated that large language models (LLMs) can automatically decompose and execute complex scientific workflows^11,12^, translating this success to the highly specialized domain of S2F modeling reveals critical limitations. General-purpose LLMs are not intrinsically grounded in the execution constraints of heterogeneous S2F systems, leaving them prone to selecting incompatible models, inferring unsupported parameters, and generating non-executable workflows. These failures arise not simply from insufficient biological knowledge, but from the absence of explicit computational and evidentiary constraints linking natural-language intent to valid scientific execution. Overcoming the fragility of general-purpose LLMs requires a structural shift toward harness engineering, a systemic computational scaffolding designed to strictly constrain agent behavior within verifiable computational and biological contracts, such as explicit input, output, routing, and evidence contracts. Within a skill-grounded harness, open-ended biological requests can be securely anchored to verifiable execution paths, effectively translating general LLMs capabilities into robust and reliable S2F workflows.

Here, we present S2F-Agent, a human-in-the-loop agent that enables verifiable orchestration of the heterogeneous S2F ecosystem. At the core, the framework explicitly decouples model-specific capabilities (Skills) from model-agnostic biological objectives (Playbooks). The harness translates free-form biological requests into constrained execution plans, providing the execution constraints that connect model-specific Skills with higher-level Playbooks. Benchmarking across routing, groundedness, and task execution showed the harness engineering of S2F-Agent could successfully reduce routing failures and analytical hallucinations inherent to general LLMs, with substantially higher accuracy across all evaluation suites. Crucially, by integrating model predictions with independent evidence, the agent gates downstream interpretation according to the strength of orthogonal genetic and phenotypic evidence, maximizing functional hypothesis generation while preventing unsupported scientific claims.

We demonstrate its utility of this framework across four levels of genomic analysis. First, we show that S2F-Agent autonomously orchestrates the adaptation of the genomic foundation model to quantitative chromatin profiles, disentangling the distinct motif signatures governing primed versus active chromatin states. Second, by harmonizing multi-perspective variant effect predictions (VEP) with quantitative genetics, we efficiently distilled CAD-associated variants into 42 high-priority regulatory candidates, recovering tissue-resolved therapeutic targets such as the hepatic SORT1 metabolic axis. Third, genome-scale profiling over 250,000 variants across multiple traits GWAS atlas reveals pervasive context dependence in molecular consequences and regulatory architecture, highlighting the analytical focus shifting from universal functional scores toward fine-grained, tissue-specific regulatory cascades. Finally, we introduce an evidence-gated analysis of personal genome interpretation that expands the functional hypothesis space beyond known clinical registries to thousands of highly prioritized candidates, maximizing discovery while imposing an evidence-dependent ceiling on downstream claims. Together, these results establish S2F-Agent as a general framework for converting heterogeneous sequence-to-function capabilities into composable, verifiable, and scalable genomic analyses.

## Results

### Overview of S2F-Agent

S2F-Agent is a human-in-the-loop agentic framework designed to autonomously orchestrate sequence-to-function (S2F) analyses for genomic interpretation through the integration of domain-specific models. Powered by Large Language Models (LLMs), the system allows users to seamlessly issue instructions to interpret genomic sequences via an intuitive natural language interface. In response, S2F-Agent automatically orchestrates sophisticated analytical pipelines, encompassing tasks such as sequence embedding, model fine-tuning, and variant effect prediction **(Figure 1A)**.

**Figure 1.**
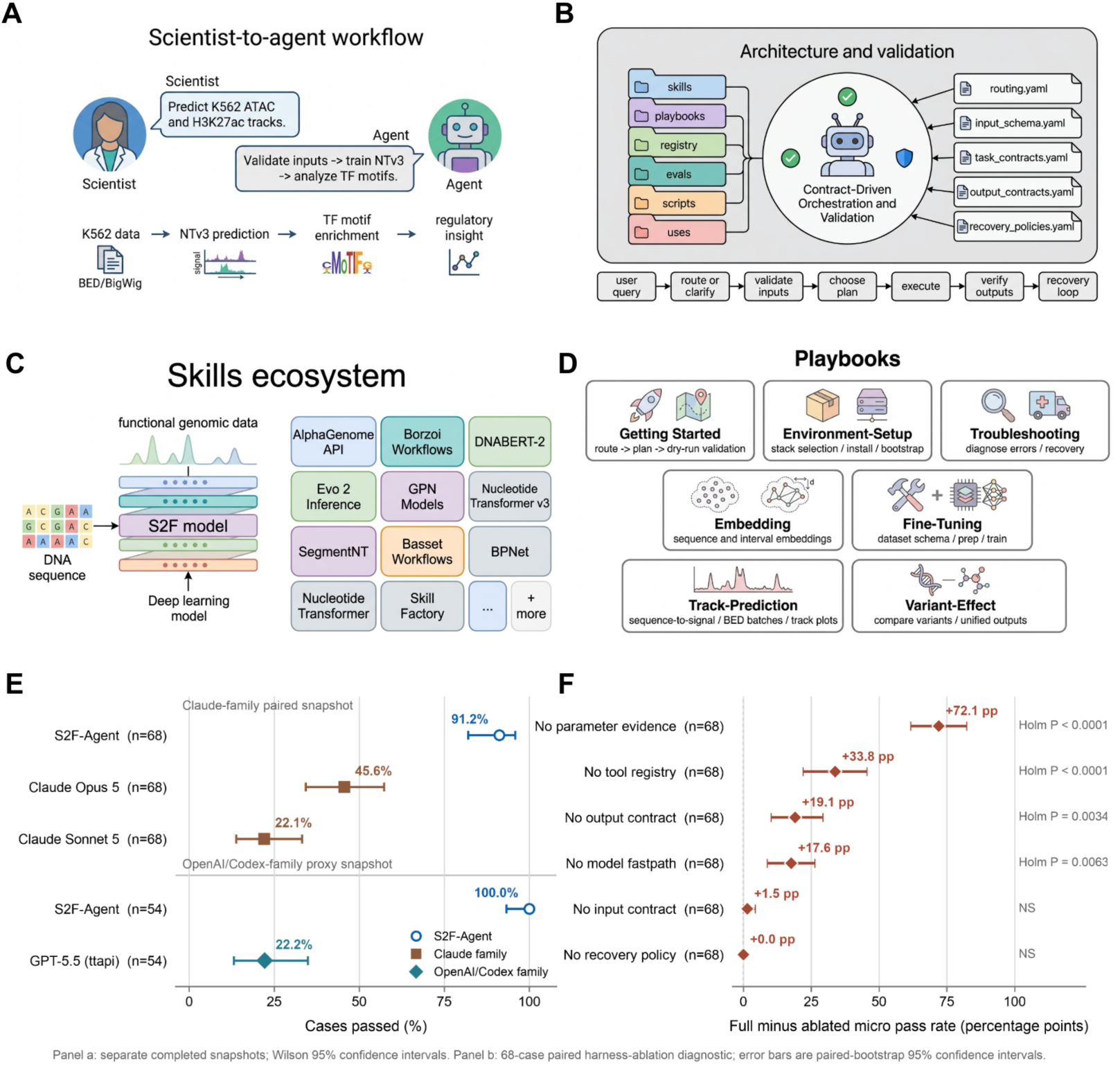
Overview of S2F-Agent as a contract-driven agent system for sequence-to-function model workflows. The figure summarizes the major components and operating logic of S2F-Agent, including the Scientist-to-agent workflow (**A**), Agent Architecture (**B**), Skills ecosystem (**C**), and Playbooks (**D**). **(A)** The Scientist-to-agent workflow panel illustrates a representative scenario in which the scientist asks a task, and the agent validates genomic inputs, selects a sequence-to-function model, runs the analysis pipeline, and returns biological insight. **(B)** The Architecture and validation panel depicts the central agent as a contract-driven orchestrator that connects repository modules, routing logic, input schemas, task and output contracts, and recovery policies. The bottom workflow indicates how user queries are routed or clarified, validated, planned, executed, verified, and recovered when needed. **(C)** The Skills ecosystem panel illustrates how sequence-to-function models transform DNA sequence inputs into functional genomic outputs, alongside representative modular skills exposed by the system, including AlphaGenome API, Borzoi Workflows, DNABERT-2, Evo 2, GPN, Nucleotide Transformer, SegmentNT, Basset, BPNet, and Skill Factory. **(D)** The Playbooks panel shows standardized workflow controllers for common biological tasks, including getting started, environment setup, troubleshooting, embedding, fine-tuning, track prediction, and variant-effect analysis. **(E)** Overall pass rates for S2F-Agent compared to generalist base models. The evaluation is divided into two separate comparisons: a 68-case paired snapshot comparing S2F-Agent (91.2%) against Claude Opus 5 (45.6%) and Claude Sonnet 5 (22.1%), and a 54-case paired snapshot comparing S2F-Agent (100.0%) against a GPT-5.5 (22.2%). Error bars represent Wilson 95% confidence intervals. System comparisons are only interpretable within each respective paired snapshot. Markers denote S2F-Agent (blue circles), Claude-family models (brown squares), and the OpenAI/Codex-family proxy (teal diamond). **(F)** Local harness ablation by paired micro pass-rate difference (full-harness advantage) between the complete S2F-Agent architecture and six leave-one-out runtime variants. The ablation highlights the critical operational roles of parameter evidence (+72.1 percentage points) and tool registries (+33.8 percentage points) in maintaining reliable tool orchestration. Error bars indicate paired-bootstrap 95% confidence intervals. Reported P values are derived from Holm-adjusted exact McNemar tests.

At the core of S2F-Agent lies a rigorous harness architecture that translates free-form user intent into the reliable execution of analytical workflows **(Figure 1B)**. By leveraging structured input and output schemas, the system normalizes ambiguous user queries into stringent contracts. A dedicated router subsequently enforces these constraints to synthesize deterministic and reproducible execution plans. Crucially, this contract protocol features a human-in-the-loop safeguard mechanism, systematically halting execution to request user clarification when biological parameters lack precise definition.

To guarantee reliable execution across a diverse repertoire of specialized sequence-to-function (S2F) models, S2F-Agent decouples specific model capabilities (Skills) from generalized biological tasks (Playbooks) **(Figure 1C, D)**. This dual abstraction ensures that model-specific syntax remains isolated, while a shared orchestration layer seamlessly coordinates complex workflows. For instance, when a user requests a variant-effect analysis, the playbook standardizes the overarching biological workflow, while the internal router dynamically dispatches the task to the optimal skill. Consequently, this modular architecture establishes S2F-Agent as a highly reusable and scalable framework for versatile cross-model orchestration.

### Harness engineering of the S2F ecosystem via Skills and Playbooks

To comprehensively cover the state-of-the-art sequence-to-function ecosystem, S2F-Agent currently registered more than 20 model-level skills (**Figure 1C**). Mirroring the operational diversity of the expanding ecosystem, these registered skills bifurcate into two primary paradigms. The first comprises direct predictive models (e.g. AlphaGenome^4^, Borzoi^5^, Enformer, ChromBPNet) that map sequences to molecular phenotypes, resolving chromatin accessibility, gene transcription, splicing, and 3D genome features. The second encompasses foundation models (e.g. DNABERT^7^, Evo^8^, GPN^9^, and NT^10^) trained on unlabeled genome-scale corpora to extract underlying representations and evolutionary constraints from genomic sequences. By embedding domain-specific expert knowledge into the agent, these integrated skills guarantee the highly reliable and standardized execution of their corresponding S2F models across both modeling paradigms.

At the orchestration layer, Playbooks bridge the semantic gap between natural user intent and backend execution capabilities (**Figure 1D**). In genomic research scenarios, these intents typically revolve around critical S2F applications, such as predicting molecular phenotypes from raw sequences, fine-tuning foundation models for domain-specific tasks, and quantifying the regulatory impact of genetic variants. Driven by the need to execute these high-level biological objectives rather than manage model-specific commands, S2F-Agent utilizes Playbooks to dynamically orchestrate underlying *Skills* into four robust workflow classes:

- Track Prediction: Normalizes requirements for species, genome assembly, sequence intervals, and output heads to structure multi-track genomic predictions, enabling the systematic inference of diverse functional readouts, such as chromatin accessibility and gene expression.
- Embedding Generation: Requires explicit sequence boundaries and embedding targets to differentiate between specific representation strategies, facilitating the extraction of the complex representation and evolutionary constraints underlying sequence.
- Fine-Tuning: Aligns training pipelines with expected artifacts by mandating dataset schemas, task objectives, and compute constraints, allowing foundation models to capture cell-type-specific regulatory dynamics within user-provided dataset.
- Variant-Effect Prediction: Enforces strict programmatic contracts for assembly, genomic coordinate, and allele specification before generating deterministic execution steps for in silico mutagenesis, thereby predicting the impact of arbitrary genetic variations on functional outputs.

### Benchmarking S2F-Agent with General LLMs

The fidelity of this harness architecture is validated by a contract-based evaluation suite. Rather than chasing generalized conversational leaderboards, the benchmark evaluates whether a system can select compatible skills, preserve grounded constraints, and generate executable plans. We evaluated 54 cases to evaluate the biological drift and tool-use success in sequence-to-function workflows, where each query in the benchmark is derived from a published study (**Supplementary Materials**). The benchmark aggregates three strict evaluation modules: a routing suite (23 cases) testing stable skill selection and ambiguity handling; a groundedness suite (8 cases) monitoring against hallucination-like fabrications or unsupported API behaviors; and a task-success suite (23 cases) verifying that generated plans are structurally complete and executable. Furthermore, to isolate the causal contributions of the system’s explicit controls, we designed a leave-one-out harness ablation. This diagnostic protocol compared the full local harness against six runtime variants that systematically removed parameter evidence, tool registries, input contracts, output contracts, recovery policies, or model-specific fastpaths.

We evaluated the S2F-Agent (based on OpenAI GPT-5.5) alongside several base LLMs, such as OpenAI GPT-5.5 and Claude Opus5, Sonnet5, reporting micro-averaged accuracies across all benchmark queries (**Figure 1E**). The results demonstrate that S2F-Agent achieved substantially higher accuracy than general LLMs across all evaluation suites, where general LLMs frequently struggle with tool orchestration, leading to analytical hallucinations and execution failures in specialized S2F scenarios. Specifically, S2F-Agent achieved a 100.00% pass rate (54 of 54 cases), whereas the GPT-5.5 passed only 22.22% (12 of 54 cases), demonstrating a significant paired overall difference of 77.78 percentage points (*P* = 4.55 × 10^−13^). Similarly, in a separate 68-case Claude-family snapshot, S2F-Agent passed 91.18% of strict cases (62 of 68). By comparison, the Claude Opus 5 and Claude Sonnet 5 compatibility conditions achieved pass rates of only 45.59% (31 of 68) and 22.06% (15 of 68), respectively. S2F-Agent outperformed Claude Opus 5 by 45.59 percentage points (*P* = 9.31 × 10^−14^) and Claude Sonnet 5 by 69.12 percentage points (*P* = 1.42 × 10^−14^) (**Supplementary Materials**). These paired outcomes collectively demonstrate that S2F-Agent achieves substantially higher accuracy in sequence-to-function tool orchestration, avoiding the execution failures and fabrications frequently observed when relying on general LLMs.

The harness ablation provided clear evidence that explicit model-specific operational knowledge drives contract-level success (**Figure 1F**). The full harness passed 95.59% (65 of 68) of the strict ablation cases. Removing structured parameter evidence caused the most severe performance collapse, dropping the overall pass rate to 23.53% (a paired loss of 72.06 percentage points; Holm-adjusted exact McNemar *P* = 6.39 × 10^−14^). Removing the tool registry also significantly degraded performance, reducing the overall pass rate to 61.8% (*P* = 3.81 × 10^−6^). Furthermore, ablating output contracts or model-specific fastpaths heavily impacted task success, passing only 23.1% and 30.8% of task-success cases, respectively. These patterns confirm that parameter evidence, tool metadata, output contracts, and fastpaths each make measurable, operationally consequential contributions to reliable sequence-to-function orchestration.

Taken together, the architectural design of S2F-Agent, anchored by the dual abstraction of Skills and Playbooks, systematically bridges the semantic gap between open-ended biological intent and rigid computational execution. By strictly constraining LLM reasoning within verifiable programmatic contracts, this harness engineering approach eliminates the hallucinations and routing failures that plague general-purpose models. Consequently, S2F-Agent established a robust orchestration engine that transforms raw linguistic reasoning into specialized S2F modeling, further primed to accelerate real-world genomic discoveries.

### Decoding regulatory grammar through foundational model fine-tuning on functional profiles

To demonstrate the capability of S2F-Agent in orchestrating sequence-to-function fine-tuning workflows, we constructed a multitask regression benchmark using ENCODE K562 ATAC-seq and H3K27ac ChIP-seq signals. Specifically, we generated a balanced dataset of 64,198 regions by anchoring fixed 4,096-bp sequence windows across active regulatory domains and randomly sampled, non-overlapping background loci. To ensure rigorous spatial generalization, we isolated chromosome 8 as a strict holdout test set (n = 2,520), splitting the remaining regions into a training set (n = 52,395) and a validation set (n = 9,283). The extracted GRCh38 reference sequences were then paired with quantitative epigenetic targets, defined as log1p-transformed and standardized mean fold-change ATAC-seq and H3K27ac signals.

Deploying the structured Nucleotide Transformer skill and fine-tuning playbook, S2F-Agent autonomously executed the lightweight adaptation of the NTv3 model on the regulatory dataset. The pre-trained backbone (InstaDeepAI/NTv3_100M_post) was strictly frozen, and a two-output linear regression head was trained on the attention-pooled 768-dimensional embeddings, to predict the quantitative chromatin states. The agent seamlessly managed the training loop with early stopping, continuously monitoring validation loss until reaching an optimal minimum (normalized MSE = 0.07019) on chromosome 7 at epoch 19 **(Figure 2a)**. Evaluated on the held-out chromosome 8 test set, the fine-tuned model demonstrated strong locus-level predictive accuracy, achieving Pearson correlation coefficients (PCC) of 0.811 (*R*^2^ = 0.653) for ATAC-seq and 0.799 (*R*^2^ = 0.637) for H3K27ac task **(Figure 2b, and Supplementary Figure 2b)**, with normalized MAEs of 0.210 and 0.211, respectively **(Supplementary Figure 2c)**.

**Figure 2.**
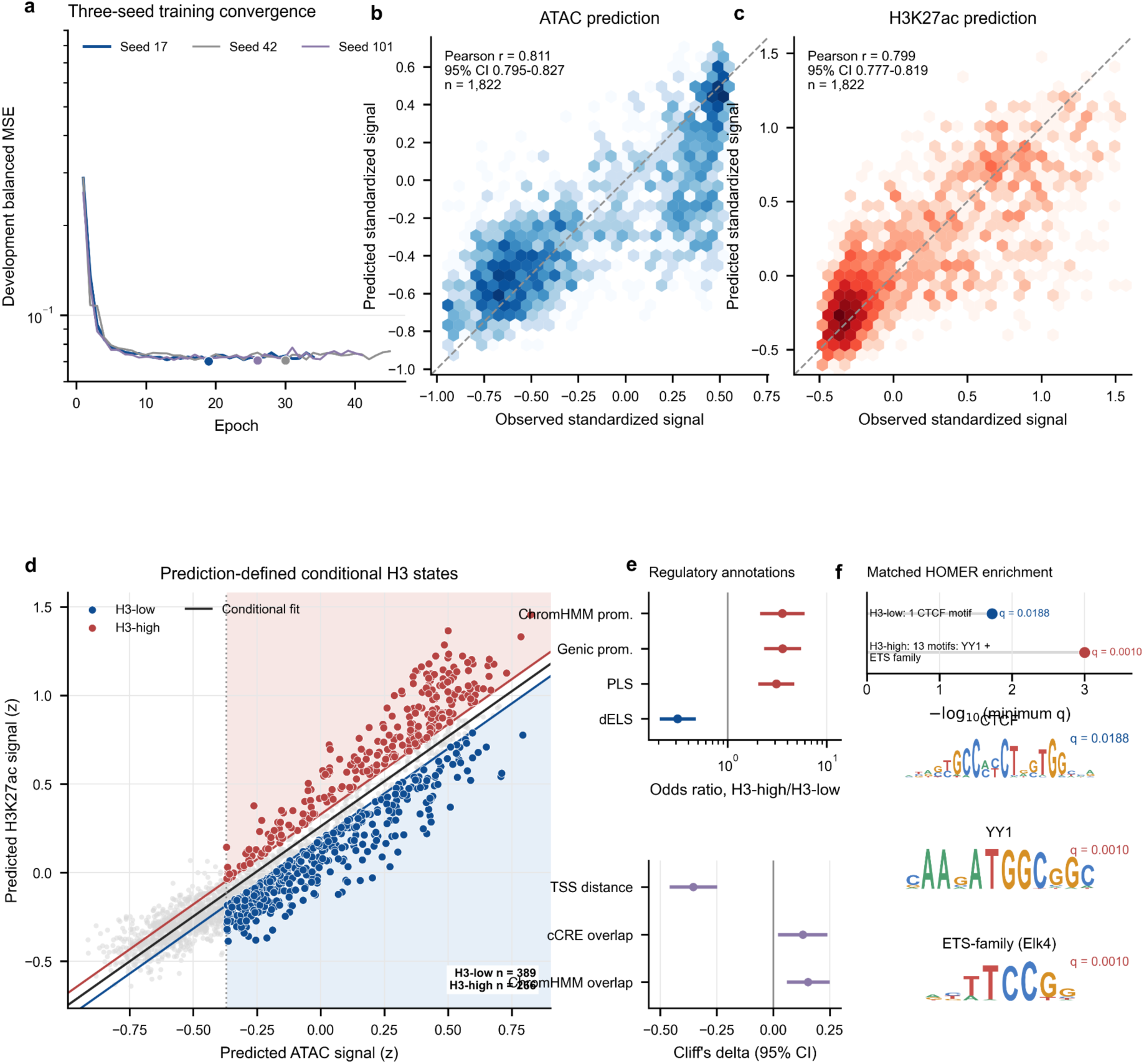
Sequence-based prediction and conditional selection of differential K562 regulatory loci. **(A)** Training and development balanced normalized mean-squared error for three frozen-encoder linear heads. The highlighted point marks the selected seed 17 checkpoint at epoch 19 (development MSE, 0.0702). **(B-C)** Locus-median observed and predicted standardized ATAC-seq (**B**) and H3K27ac (**C**) signals for 1,822 held-out chr8 loci. Hexagons encode locus density; dashed lines denote identity and solid lines denote fitted calibration. R2 values and percentile 95% confidence intervals were estimated by 2,000 bootstrap resamples of loci. **(D)** Conditional H3K27ac-versus-ATAC landscape used to identify H3-low/ATAC-high loci. The conditional fit and thresholds were defined on chr7 development data. Dark blue points are 94 loci meeting the strict Q10 residual/Q60 ATAC rule; light-blue points add 99 loci under the post-hoc relaxed Q20/Q50 rule (193 total). **(E)** Agreement of three prediction heads with the observed-signal relaxed candidates. Circles show agreement with the negative conditional-residual direction; squares show agreement with the complete relaxed rule. Candidate membership is defined only from observed signals. **(F)** Exploratory motif analysis of the 193 relaxed candidates against 579 non-reused matched background loci. AME tested 1,019 JASPAR2026 motifs and reported no motifs at q <= 0.1. Standardized mean differences summarize matching on GC fraction, CpG density, joint signal and locus length. The relaxed analysis is post-hoc and lacks mappability, TSS-distance and ChromHMM matching and orthogonal K562 binding validation; motif results are therefore exploratory.

To elucidate the biological insights captured by the fine-tuned model, we examined the sequence grammar governing the epigenetic heterogeneity between ATAC-seq signals and H3K27ac marks. Based on the discordant relationship of these predicted profiles across the held-out chromosome, we identified two divergent regulatory classes: H3K27ac-low/ATAC-high and H3K27ac-high/ATAC-high, representing primed and active regulatory states within accessible chromatin **(Figure 2d, and Supplementary Figure 2d,e)**. Genomic enrichment analyses revealed that H3K27ac-high loci are predominantly anchored in promoter architecture, whereas H3K27ac-low/ATAC-high loci primarily represent distal regulatory domains. Specifically, H3K27ac-high regions exhibited striking proximity to transcription start sites (median absolute TSS distance of 87 bp vs. 6,128 bp; Cliff’s *δ* = −0.405, *q* = 2.05 × 10^−15^), a higher prevalence of promoter ChromHMM states (88.2% vs. 51.1%; odds ratio [OR] = 7.11, 95% CI: 4.65–10.85), and a marked enrichment of promoter-like candidate cis-regulatory elements (PLS; 54.8% vs. 21.5%; OR = 4.40). Conversely, distal enhancer-like elements (dELS) were heavily depleted relative to H3K27ac-low/ATAC-high regions (36.2% vs. 68.2%; OR = 0.266) **(Figure 2c, and Supplementary Figure 2f)**. Transcription factor bind site (TFBS) enrichment analysis further delineated the motif signatures underlying these distinct regulatory classes. Following a covariate-matched sensitivity analysis (controlling for GC content, CpG density, ATAC signal, peak intensity, and locus length), the H3K27ac-high regions exhibited significant enrichment for ETS1 and YY1 motifs, consistent with their established roles in driving promoter activity and coordinating active transcriptional machinery. Conversely, the H3K27ac-low regions were uniquely enriched for CTCF motifs, suggesting that these accessible yet hypoacetylated regions primarily function as structural chromatin boundaries or insulated regulatory hubs **(Figure 2d)**.

In summary, these results demonstrate that automated fine-tuning orchestration via S2F-Agent effectively disentangles complex multi-modal epigenomic states, translating sequence-to-function predictions into a highly interpretable regulatory principles such as the divergence between primed and active chromatin states.

### Multi-Perspective Variant Effect Prediction Decodes Non-Coding Pathogenicity in Complex Diseases

Recent advances in sequence-to-function (S2F) deep learning have revolutionized variant effect prediction (VEP) by directly mapping non-coding sequence variation to specific molecular function perturbation. However, accurately decoding non-coding pathogenicity in complex diseases remains challenging, as a disease-causing mutation perturb complex regulatory architecture, including tissue-specific chromatin accessibility, disrupt gene expression, or violate deep evolutionary constraints. Here, we utilized S2F-Agent to automatically provide a multi-perspective VEP approach that harmonizes these orthogonal functional predictions with quantitative genetic evidence for comprehensive pathogenicity prioritization in complex diseases **(Figure 3a)**.

**Figure 3.**
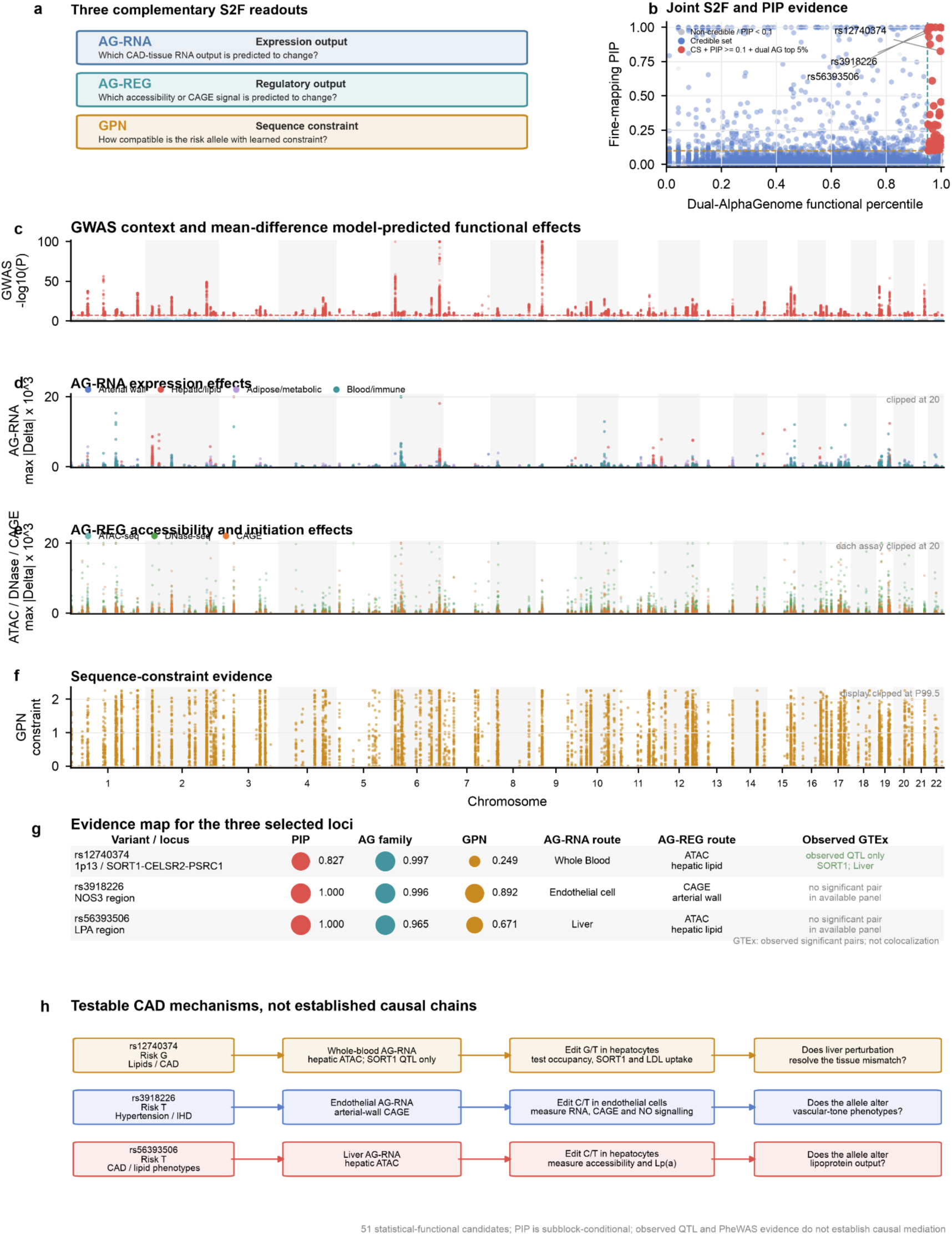
Multi-perspective S2F predictions and quantitative-genetic evidence nominate prioritized CAD mechanisms. **(a)** Three sequence-to-function (S2F) perspectives: AG-RNA predicts expression output in four prespecified CAD contexts, AG-REG predicts accessibility and transcription-initiation output in four CAD tissue groups, and GPN measures tissue-agnostic sequence constraint. **(b)** Dual-AlphaGenome functional percentile versus fine-mapping posterior inclusion probability (PIP) for 16,396 variants with matched PIP. The dual percentile is the minimum of cohort-ranked AG-RNA magnitude and the strongest assay-specific AG-REG percentile, requiring concordant ranking of the correlated AlphaGenome outputs. Grey, blue and red denote non-credible-set variants, credible-set members and the operational priority set, respectively; dashed lines mark percentile = 0.95 and PIP = 0.1. The 51 red variants meet credible-set membership, PIP >= 0.1 and dual-AlphaGenome top-5% criteria. **(c)** CAD GWAS context with 16,607 genome-wide-significant, S2F-scored variants overlaid in red; the dashed line marks P = 5 x 10^-8 and -log10(P) is clipped at 100. **(d)** Maximum absolute risk-aligned AG-RNA mean(ALT - REF) across the four CAD contexts, coloured by the maximizing context, multiplied by 10^3 and clipped at 20. **(e)** Corresponding maxima for ATAC-seq, DNase-seq and CAGE across CAD tissue groups; each head is multiplied by 10^3 and clipped at 20 without pooling raw scales. **(f)** GPN constraint strength, max(0, -risk-allele LLR), clipped at its 99.5th percentile. **(g)** PIP, dual-AlphaGenome and GPN percentiles, dominant AG-RNA context, strongest ATAC/DNase/CAGE route and observed GTEx significant-pair evidence for rs12740374, rs3918226 and rs56393506; point area and printed values encode evidence magnitude. **(h)** Risk-aligned phenotype associations and model-predicted tissue routes motivate isogenic perturbation tests. PIP is conditional on the original EUR LD-aware subblock; observed GTEx pairs are not colocalization, and no evidence layer is interpreted as a causal probability.

Leveraging our standardized orchestration framework, we sought to elevate functional variant prioritization in realistic complex traits, using coronary artery disease (CAD) as a model system. We constructed a comprehensive functional prediction manifest of 16,607 genome-wide significant allele-oriented CAD variants (*P* ≤ 5 × 10^−8^). To synthesize predictive signals, we evaluated these variants across three distinct biological paradigms: Expression output via AlphaGenome RNA predictions (AG-RNA) across four CAD-relevant cell types/tissues (endothelial cells, whole blood, liver, and visceral adipose tissue). Epigenomic regulation via AlphaGenome regulatory heads (AG-REG) spanning five assays (ATAC-seq, DNase-seq, CAGE, TF ChIP, and Histone ChIP) across four CAD tissue groups. Evolutionary sequence constraint measured by GPN log-likelihood ratios (**Figure 3c-f, and Supplementary Figure 3a)**. In parallel with functional scoring, we performed European LD-aware fine-mapping using SuSiE-RSS across 191 parent loci, identifying 3,002 credible-set variants and 605 variants with posterior inclusion probability (PIP) ≥ 0.1. To overcome the limitations of single-model scoring or unconstrained statistical prioritization, we defined an operational statistical-functional priority set requiring credible-set membership, *PIP* ≥ 0.1, and a dual-AlphaGenome functional percentile ≥ 0.95. This strict tripartite intersection successfully narrowed the broad CAD landscape into 42 high-priority statistical-functional candidate variants (**Figure 3b**).

To independently contextualize these candidates within human disease genetics, S2F-Agent automated allele-aligned cross-referencing with GTEx v8 eQTL summaries across CAD-relevant tissues and UKB-TOPMed phenome-wide association (PheWAS) profiles. In total, 33 out of 42 priority candidates harbored at least one significant allele-aligned eQTL in CAD-relevant tissues. The multi-perspective convergence is clearly illustrated by the top candidate, rs12740374 at 1p13 (*PIP* = 0.827). Functional predictions highlighted rs12740374 as driving strong whole-blood expression perturbation alongside the most pronounced reduction in hepatic-lipid chromatin accessibility (risk-aligned delta = −0.112). In agreement with the predicted hepatic footprint, where the CAD risk G allele exhibits a striking association with lipoid metabolic disorders in UKB-TOPMed PheWAS (*P =* 6.9 × 10^−53^, risk-aligned *β* = 0.160). The clinical phenotype is also underscored mechanistically by GTEx eQTL evidence connecting the risk allele to decreased hepatic SORT1 expression (normalized effect size = −1.273), establishing a cohesive biological hypothesis wherein non-coding chromatin perturbation suppresses hepatic SORT1 to defective LDL clearance and accelerate CAD pathogenesis (**Figure 3g,h, and Supplementary Figure 3b-d**). This multi-perspective capacity extends seamlessly across diverse vascular and metabolic loci. For instance, rs3918226 (NOS3 locus) pairs predicted endothelial transcriptional disruption with a pronounced PheWAS signal for hypertension (*P* = 1.2 × 10^−24^). Similarly, at the LPA locus, rs56393506 links predicted hepatic ATAC-seq attenuation directly to coronary atherosclerosis risk (*P* = 1.6 × 10^−51^) (**Figure 3g,h, and Supplementary Figure 3b-d**).

Taken together, these findings demonstrate the utility of S2F-Agent in deciphering the complex functional architecture of human disease genetics. By orchestrating multi-perspective sequence-to-function models alongside quantitative genetics, the agent effectively transforms thousands of CAD candidate variants into a transparent, tissue-resolved roadmap for targeted experimental validation and therapeutic discovery.

### Multimodal Variant Effect Profiling Delineate Context-aware Cross-trait Genetic Architecture

Although genome-wide association studies (GWAS) have identified millions of trait-associated loci, the role of context-specific variant effects in shaping overarching cross-trait genetic architecture remains a major unresolved question. To establish context-aware functional priorities for cross-trait genetic landscape, we deployed the S2F-Agent framework to automatically perform large-scale variant effect profiling across the ontology-annotated GWAS Catalog dataset, comprising thousands of association records normalized through standardized ontology crosswalks. Specifically, we reorganized heterogeneous association records into a hierarchical phenotype specification that structured fine-grained entities across four major disease anchors, including coronary artery disease (CAD), type 2 diabetes (T2D), chronic kidney disease (CKD), and metabolic dysfunction-associated steatotic liver disease (MASLD). Then, we explicitly separated core disease anchors from clinically nested phenotypes, such as CAD with myocardial infarction, and CKD with eGFR **(Supplementary Figure 4a,b)**. In total, we mapped 15 fine-grained phenotype entities, 4 disease anchors, and 250,287 model-compatible single-nucleotide variants (SNVs) into three distinct S2F prediction perspectives, including AlphaGenome expression (AG-RNA), AlphaGenome regulation (AG-REG), and GPN sequence constraint.

Across the predicted functional atlas of genome-wide variants, these multi-perspective modalities capture complementary, non-redundant molecular mechanisms and evolutionary conservation rather than a single monolithic priority score. Evaluation of task correlations across the predicted atlas demonstrated that AlphaGenome expression and regulation outputs were moderately correlated (Spearman *ρ* = 0.441, chromosome-cluster bootstrap 95% confidence interval: 0.426–0.449) **(Figure 4a)**. In contrast, GPN sequence constraint was essentially orthogonal to both expression (*ρ* = −0.0076) and AG-REG (*ρ* = 0.0105), indicating that GPN captures evolutionary sequence properties distinct from AlphaGenome molecular perturbations. Within AG-REG, correlations among ATAC, DNase, CAGE, histone ChIP and transcription-factor ChIP aggregates ranged from 0.468 to 0.851, indicating related but non-interchangeable regulatory outputs **(Supplementary Figure 4c,d)**. To identify functionally significant loci, variants were evaluated against a fixed atlas-wide 95th-percentile (P95) threshold across all three prediction axes. Single-axis support predominated, comprising 31,475 AG-RNA-only, 7,385 AG-REG-only, and 9,843 GPN-only variants, whereas top-tier multi-modal concordance reaching P95 across all three S2F axes simultaneously was observed in only 550 variants (0.22%) **(Supplementary Figure 4e)**.

**Figure 4.**
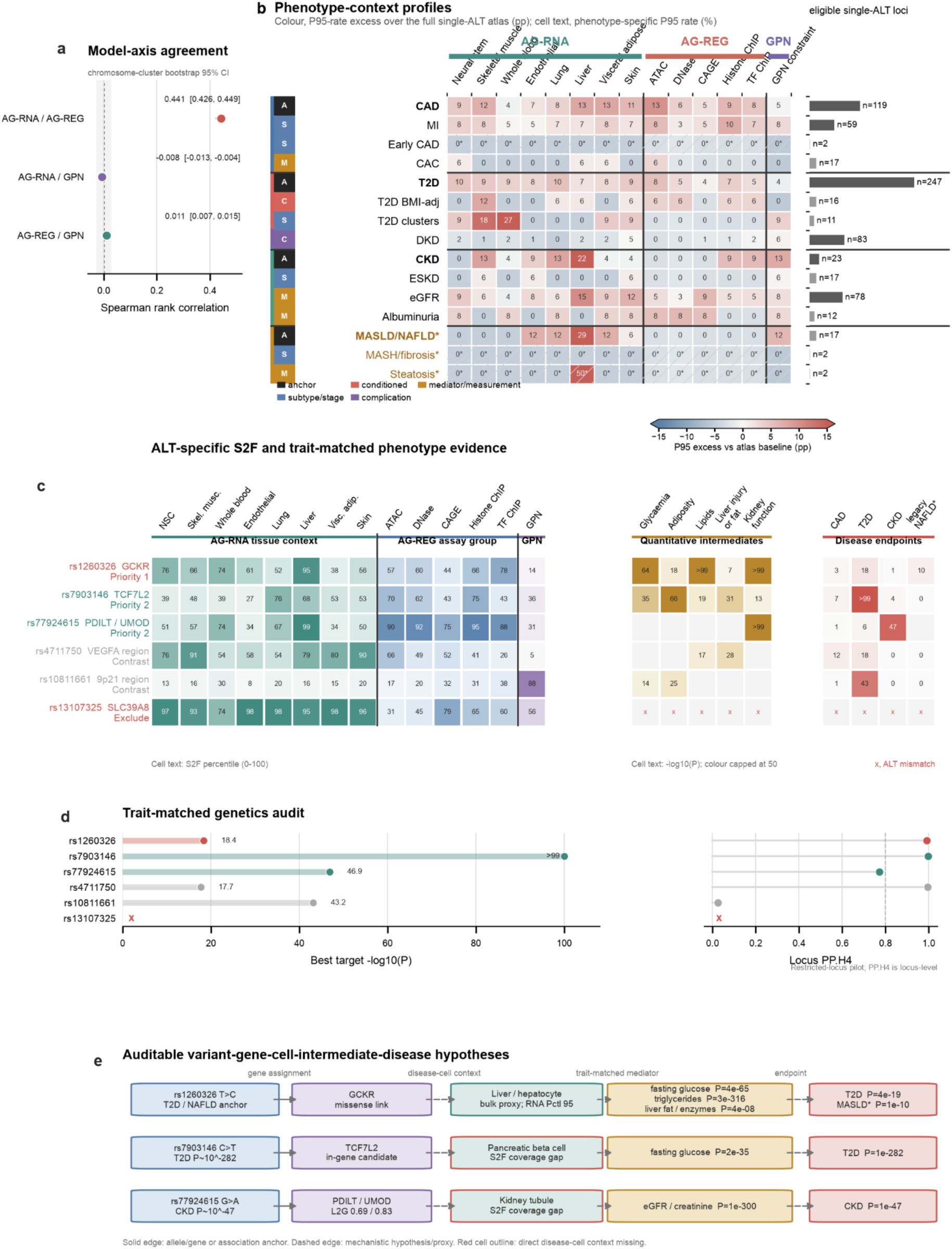
Contract-aware variant-effect profiles define testable cross-trait hypotheses. **(a)** Pairwise Spearman rank correlations between the mean AG-RNA, AG-REG and GPN percentile axes for 250,287 ALT-specific variants. Points denote Spearman rho and horizontal lines denote 95% confidence intervals from 1,000 chromosome-cluster bootstrap resamples; AG-RNA and AG-REG are correlated tasks from one model family and are not independent replication. **(b)** Phenotype-context profiles for 15 entities frozen before joining model outputs. Columns comprise eight AG-RNA tissue or cell contexts, five AG-REG assay-group aggregates and GPN. Cell text gives the percentage of eligible single-ALT reported loci at or above the axis-specific P95 threshold; colour gives the percentage-point difference from the corresponding empirical single-ALT atlas baseline (n = 161,553 rsIDs), and right-hand bars give eligible-locus counts. Hatching marks n < 10. Multi-ALT rsIDs are excluded. **(c)** Six prespecified cases ordered by actionability tier. S2F cells give within-axis percentiles (0–100); quantitative-intermediate and disease-endpoint cells give -log10(P), with colour capped at 50. An x denotes an ALT mismatch that precludes integration. **(d)** Restricted ten-locus genetics audit. Left, strongest target-disease association as -log10(P); right, maximum main-prior PP.H4 for a locus-level target-trait pair. The dashed line marks 0.8; PP.H4 is locus-level rather than variant-specific. **(e)** Auditable hypotheses for rs1260326, rs7903146 and rs77924615. Blue, purple, teal, gold and red nodes denote variant, candidate gene, cell or tissue, quantitative intermediate and disease endpoint, respectively. Solid arrows mark allele, gene or association anchors; dashed arrows mark unvalidated mechanistic links, and red cell outlines mark absent direct beta-cell or kidney-tubule coverage. These profiles are descriptive and do not establish enrichment, shared causality, effector genes or therapeutic targets; confirmatory Tier B remains STOP.

**Figure 5.**
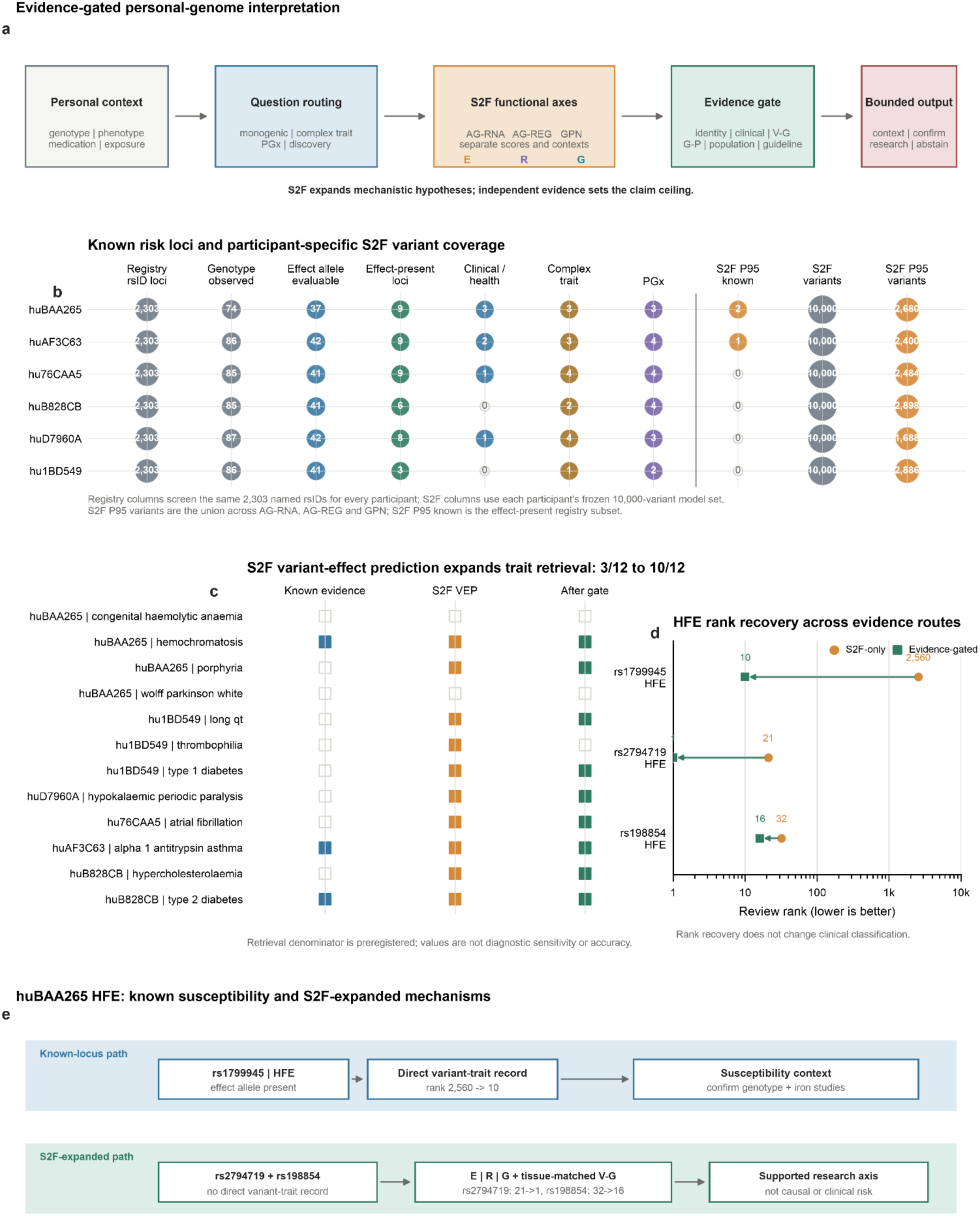
Layered evidence for personal-genome interpretation. a, Question-routed workflow linking participant genotype and phenotype context, separate AG-RNA, AG-REG and GPN outputs, independent evidence and bounded reporting. b, Complete-registry screen of six participants. Each receives the same 2,303 named rsID loci; 74-87 are genotype-observed, 37-42 have an evaluable registry effect allele and 3-9 are effect-present. Clinical/health, complex-trait and PGx classes partition effect-present loci; S2F P95 is a functional subset. c, Retrieval across 12 preregistered self-reported phenotype groups. Known evidence retrieves 3/12 groups, S2F variant-effect candidates retrieve 10/12, a gain of seven groups, and 9/12 retain a lead after the available evidence gate. Retrieval is a contextual endpoint, not diagnostic sensitivity or independently validated prediction accuracy. d, Review-rank change for three known loci in the frozen S2F input. Circles denote S2F-only rank and squares denote evidence-gated rank; lower ranks indicate earlier review without changing source classification. e, huBAA265 HFE comparison between known rs1799945 susceptibility context and S2F-expanded rs2794719 and rs198854 mechanisms. Expanded paths remain supported research axes rather than causal or personal-risk estimates.

Evaluating the fine-grained phenotype entities across s2f-model predictions revealed distinct context-specific functional effects of variants, instead of a universal cross-disease effect **(Figure 4b)**. Specifically, bulk liver AG-RNA emerged as a primary organ-level context shared across hepatic and renal entities, led by MASLD/NAFLD (+24.6 percentage points [pp] excess over the atlas baseline; n = 17) and hepatic steatosis (50% P95 rate), alongside chronic kidney disease (CKD; +16.9 pp excess, n = 23) and its quantitative intermediate eGFR (15% P95 rate, n = 78). For the regulatory modalities, Coronary artery disease (CAD) and myocardial infarction (MI) exhibited their primary functional footprints in chromatin accessibility (+ 9.9 pp excess, n = 119; and 8% P95 rate, n = 59), complemented by histone ChIP marks (11/119; 9.2%; +6.3 pp excess; and 59; 8.5%; +5.8 pp excess, respectively). The predicted profiles also reflect a fundamental trade-off between broad activity and context selectivity, where canonical disease anchors exhibit multi-modal support, whereas refined sub-phenotypes demonstrated distinct tissue specificity, including T2D clusters in whole blood (27% P95 rate, +22 pp excess) and skeletal muscle (18%), alongside BMI-adjusted T2D in skeletal muscle (12%).

To translate the predicted landscape into biological mechanisms, we generated auditable hypotheses by integrating multimodal variant effect profiles with candidate-gene annotations, trait-matched quantitative intermediates, and clinical disease endpoints **(Figure 4c)**. The most complete, experimentally actionable hypothesis emerged at rs1260326 (T>C). This variant exhibited a liver AG-RNA percentile of 0.953 and a transcription-factor ChIP regulatory percentile of 0.778, mapping directly to a coding missense variant in GCKR. Trait-matched genetic auditing linked rs1260326 to strong quantitative intermediate phenotypes—including fasting glucose (*P* = 4 × 10^−65^), serum triglycerides (*P* = 3 × 10^−316^), and liver fat/enzymes (*P* = 4 × 10^−8^)—which sequentially connected to clinical endpoints for type 2 diabetes (T2D; (*P* = 4.2 × 10^−19^) and metabolic dysfunction-associated steatotic liver disease (MASLD/legacy-NAFLD;*P* = 1.1 × 10^−10^). This coherent cascade supports a GCKR-centered, liver-mediated metabolic axis, though bulk liver predictions represent an organ-level proxy that cannot yet disentangle direct horizontal pleiotropy from vertical metabolic mediation. Complementary cases demonstrated how disease-anchored loci can prioritize intermediate trait cascades while exposing critical cellular coverage gaps. At rs7903146 (C>T), a robust T2D association (*P* = 1.1 × 10^−282^) and an in-gene candidate link to TCF7L2 aligned with a fasting glucose intermediate signal (*P* = 2 × 10^−35^). Similarly, rs77924615 (G>A) connected chronic kidney disease (CKD; (*P* = 1.4 × 10^−47^) and an eGFR/creatinine intermediate (*P* = 1 × 10^−300^) to high locus-to-gene mapping scores for PDILT (0.693) and UMOD (0.826).

Integrating quantitative genetics and population-genetic controls could further establish rigorous boundaries on these hypotheses **(Figure 4d,e)**. A restricted statistical genetic analysis across ten locked ±500-kb loci (119,476 trait-variant rows) identified 138 conditional signals and approximate 95% credible sets via reference-LD conditioning. Across 60 locus-by-trait-pair colocalization comparisons, four loci were classified as trait-specific candidates and six as unresolved, with zero shared credible candidates passing the confirmatory colocalization gate (PP.H4). Furthermore, atlas-wide population-genetic matching (129,579 exact 1000 Genomes Project matches) demonstrated that S2F percentiles were largely orthogonal to minor allele frequency and ancestry differentiation (|*ρ*| ≤ 0.018). Several negative controls illustrated key failure modes across the actionability checklist **(Figure 4d)**. For instance, rs4711750 (VEGFA region) exhibited strong genetic associations (*P* = 10^−17.7^) and extreme population differentiation (*F_ST_* = 0.481), but lacked P95 multimodal S2F support, serving as an ancestry/LD alternative-explanation control. rs10811661 (9p21 region) reached high evolutionary sequence constraint (GPN = 0.883) and T2D association (*P* = 10^−43^), but lacked AG-RNA predictions and candidate-gene links.

Together, multimodal orchestration of S2F-Agent could effectively prioritize functionally active variants across a broad cross-trait atlas, establish traceable, context-bounded hypotheses linking variants, genes, cell contexts, and intermediate traits. By decoupling functional predictions from raw reported GWAS-locus, these findings demonstrated that disease-level locus sharing is primarily governed by intermediate phenotypic structures, clinical nesting, and context-dependent molecular pathways rather than universal functional scores.

### Evidence-gated Sequence-to-Function Interpretation of Personal Genomes

While personal genomes are increasingly accessible, their interpretation remains severely bottlenecked by the extreme sparsity of established clinical evidence registries. Sequence-to-function (S2F) models offer a transformative approach to this bottleneck by predicting the distinct molecular consequences of any carried allele. However, molecular predictions are not direct probabilities of pathogenicity or disease diagnoses, and misinterpreting these model outputs dangerously obscures missing context and risks hallucinating medical claims. To resolve this tension, personal-genome orchestration demands a strict evidence-gated paradigm, which systematically routes S2F predictions through independent verification layers, including clinical variant records, quantitative genetics, molecular-trait associations, participant phenotypes, pharmacogenomic resources, and environment exposure boundaries. Accordingly, the central objective was to determine whether a unified orchestration framework could successfully scale S2F scoring and link predicted variant effects to independent gene and phenotype evidence without triggering unsupported clinical escalation.

To establish a baseline capability for personal genome interpretation, we first screened six public Personal Genome Project (PGP) participants against a comprehensive clinical registry containing 3,080 source records and 2,303 named risk loci (rsIDs). This complete-registry screen starkly illustrated the inherent sparsity of known clinical evidence. For any given participant, only 74 to 87 of these registered loci were directly observed in their genotype, yielding a mere 3 to 9 verified effect-present alleles per individual. In aggregate, this screening recovered 44 effect-present observations across 19 rsIDs, which were thinly scattered among clinical, complex-trait, and pharmacogenomic classes. To overcome this evidence sparsity, S2F-Agent constructed a highly granular, participant-resolved functional landscape by profiling 10,000 carried alleles per participant (54,099 unique alleles globally), across AlphaGenome RNA (AG-RNA), AlphaGenome regulatory (AG-REG), and GPN (evolutionary constraint) modalities. In total, this multi-modal prioritization yielded between 1,688 and 2,898 top-percentile (P95) functional candidates per individual (exceeding 15,000 high-impact observations cohort-wide), which significantly extends the hypothesis spaces of variant candidates.

To determine whether this expanded functional landscape generates biologically meaningful signals rather than computational noise, we evaluated variant retrieval across 12 preregistered participant phenotypes based on Human Phenotype Ontology (HPO). While known clinical evidence linked variants to only 3 of the 12 groups, S2F profiling retrieved contextually relevant functional candidates for 10 groups, substantially increasing explanatory coverage. To prevent false-positive escalation, S2F-Agent further subjected these candidates to stringent independent evidence gating via variant-to-gene mapping. Under the strict restraint, 9 of the 12 phenotype groups retained a highly prioritized, evidence-bounded mechanistic lead **(Fig. 1c)**. A locus-level view of participant huD7960A exemplifies this dynamic, demonstrating how S2F signals nominated relevant functional candidates across the CACNA1S and KCNJ2 regions for hypokalaemic periodic paralysis, while strictly down-ranking unsupported variants during subsequent target-gene review **(Supplementary Fig. 3a)**. Although the endpoints represent contextual retrieval rather than diagnostic sensitivity, these results definitively demonstrate that evidence-gated S2F scoring effectively bridges the gap between abstract molecular predictions and real-world traits, capturing plausible functional mechanisms that traditional registries entirely miss.

Generating functional hypotheses is only half the computational challenge, while preventing AI models from hallucinating unverified clinical claims is the critical counterpart. To enforce this epistemic separation, S2F-Agent routed the expanded candidates through strict evidence gates across clinical variant records, quantitative genetics, and molecular-trait associations boundaries. This rigorous gating effectively distilled expansive functional signals into tightly bounded mechanistic hypotheses. For instance, among 43 uncertain candidates (24 phenotype-linked and 19 low-frequency variants), lexicographic gating successfully connected 21 variants into complete variant-effect-gene-phenotype research axes. Furthermore, a stringent independent non-coding benchmark contracted these 43 candidates down to 37 eligible non-coding variants, yielding only 4 with independent, tissue-matched variant-to-gene support. The discriminatory power of this evidence-gated logic is further illustrated by the HFE (hemochromatosis) locus in participant huBAA265. For the known risk allele rs1799945, independent variant-trait evidence correctly bounded the finding at a susceptibility-context ceiling, accelerating its review rank from 2,560 to 10. Conversely, S2F profiling uncovered two non-coding HFE candidates (rs2794719 and rs198854) that lacked direct variant-trait records in public registries. Bolstered by strong S2F effects, tissue-matched HFE eQTLs, and HPO disease links, the agent dynamically elevated their ranks from 21 to 1 and 32 to 16, respectively **(Fig. 1d)**.

Crucially, this evidence-gated framework actively expands pharmacogenomic (PGx) and gene-environment (GxE) evaluation spaces while enforcing strict clinical boundaries. The comprehensive PGx screen captured 20 effect-present marker observations, successfully expanding beyond 14 legacy-registry markers to include 6 CYP3A5 additions. This expansion matched self-reported medications not only to established clinical markers (pravastatin-SLCO1B1 and warfarin-CYP2C9), but also yielded S2F-supported research context for propafenone-CYP2D6 **(Supplementary Fig. 3c)**. However, because standard array data cannot fully resolve diplotype phase or copy-number variation, the agent rigorously abstained from inferring metabolizer phenotypes or actionable dose recommendations. GxE contextualization exhibited parallel rigor, testing 11 preregistered variant-exposure targets across a strict, multi-step evidence chain **(Supplementary Fig. 3b)**. By identifying 8 missing exposure doses or timings, 2 context mismatches, and 1 evidence conflict, the framework refused to force individualized interaction effects from mere genetic co-occurrence, systematically converting missing contexts into explicit, auditable abstentions.

Together, while measuring contextual retrieval rather than diagnostic sensitivity, these results demonstrate a profound conceptual shift in personal genome interpretation. Crucially, by rigorously relying on independent real-world evidence to dictate the claim ceiling, S2F-Agent maximally expands the molecular hypothesis space, while ensuring that an unsupported hypothesis is retained and never hallucinated as a clinical fact.

## Discussion

While sequence-to-function (S2F) models have revolutionized our ability to map non-coding genetic variation to molecular phenotypes, their explosive proliferation has shifted the central bottleneck from predictive capability to the reliable orchestration of heterogeneous models in genomic analysis. S2F-Agent addresses this challenge through a harness-based architecture that provides a unified interface between biological intent and diverse S2F capabilities. The core methodological innovation of this framework lies in the explicit decoupling of model-specific capabilities from biological objectives through Skills and Playbooks. Specifically, Skills encapsulate the execution requirements of individual S2F models, including their inputs, outputs, and operational constraints, whereas Playbooks encode higher-level biological objectives independently of any particular model implementation. This dual abstraction systematically isolates model-specific complexity, allowing newly introduced S2F models to be incorporated seamlessly without redesigning higher-level workflows. Crucially, this harness engineering effectively circumvents qualitatively distinct failure modes, including inappropriate skill selection, unsupported tool behavior, and incomplete execution plans. Evaluated against published S2F workflows, S2F-Agent substantially outperformed general-purpose LLMs, achieving superior benchmark accuracy across routing, groundedness, and task-success evaluations.

Beyond establishing computational reliability, S2F-Agent empowers systematic biological discovery across multiple scales of the sequence-to-function ecosystem. First, the fine-tuning workflow demonstrated that the framework can autonomously orchestrate model adaptation to quantitative functional profiles, effectively distilling motif interpretation from sequence representations. Second, the CAD analysis underscored that actionable variant interpretation requires the harmonization of heterogeneous S2F predictions with independent quantitative genetics, to prioritize a restricted set of regulatory candidates and generate tissue-resolved mechanistic hypotheses. Following that, genome-scale profiling across multi-trait GWAS atlas revealed a functional landscape dependent on tissue, assay, and phenotypic context, demonstrating that distinct disease entities leave tissue-specific regulatory footprints. Finally, the personal-genome analysis expands the functional hypothesis space beyond known clinical registries, constrained by a strict evidence-dependent ceiling that prevents the inflation of downstream claims. These downstream analyses highlight a complementary requirement for scientific agents: the ability to expand hypothesis generation without extending evidentiary claims beyond what available data support. By design, S2F-Agent enforces this critical distinction by generating predictions for variants absent from existing registries and integrating them with diverse genetic, molecular, and environmental evidence. This safeguard is particularly essential for individual genomes, where incomplete phenotype information, uncertain genotype phase, missing exposure data, and limited tissue coverage can otherwise encourage severe overinterpretation.

Although S2F-Agent establishes a robust foundation for automated sequence analysis, we recognize several technical boundaries. First, the system’s analytical capacity remains inherently tied to the capabilities, calibration, and biological coverage of its underlying S2F models (Skills). The orchestration layer cannot compensate for systematic model bias or inaccurate functional predictions. Second, although the benchmark provides a controlled comparison with general-purpose LLMs, it does not exhaust the space of possible genomic analyses, necessitating future prospective evaluations on independently constructed adversarial tasks. As future advancements in LLMs and S2F models will naturally alleviate the analytic bottlenecks, the modular design of S2F-Agent also guarantees a clear pathway for continuous ecological expansion. The strict decoupling of Skills and Playbooks allows the agent to seamlessly assimilate new biological modalities, while maintaining higher-level workflows within deterministic execution contracts. Consequently, S2F-Agent serves a continuously evolving ecosystem, poised to rapidly integrate emerging state-of-the-art models as the field advances.

Ultimately, S2F-Agent represents more than a natural-language interface for existing genomic tools, which marks a fundamental shift toward robust agentic orchestration. By decoupling model capabilities from biological objectives, enforcing strict execution contracts, and establishing evidence gates for downstream claims, the framework provides a general strategy for translating rapidly evolving S2F models into reproducible and scalable scientific workflows. By replacing fragmented manual analysis with a deterministic computational engine, S2F-Agent is poised to unlock the full potential of sequence-to-function ecosystems across downstream applications, including clinical genetics, therapeutic target discovery, and precision medicine.

## Data availability

Published papers used in the case studies can be found in previous studies for ENCODE: ENCFF360VIU, ENCFF849TDM; GWAS summary: GCST90132314, GCST90132315; PGP: hu1BD549, hu76CAA5, huAF3C63, huB828CB, huBAA265, huD7960A. The gwas-catalog associated datasets can be downloaded in https://www.ebi.ac.uk/gwas/. The Agent evaluation dataset is available from GitHub (https://github.com/JiaqiLiZju/s2f-agent).

## Code availability

Code for running the agent on case studies and for evaluation is available in the S2F-Agent GitHub repository (https://github.com/JiaqiLiZju/s2f-agent).

## Acknowledgments

The authors acknowledge that ChatGPT, Gemini and Nano Banana (Google) was used for assistance with language editing and drew the illustration figures of the manuscript.

## Author contributions

J.L. and Z.B. conceptualized the study. J.L. developed and implemented S2F-Agent. J.L. and J.L.(Gavin) benchmarked S2F-Agent against the base LLMs. J.L. and T.Q. collected the data, performed analysis, and evaluated results of the downstream case studies. J.L. and Z.B. supervised the study. All authors contributed to writing the paper.

## Ethics approval

Not applicable.

